## Supplementary figures for "Histone chaperone ASF1 mediates H3.3-H4 deposition in Arabidopsis"

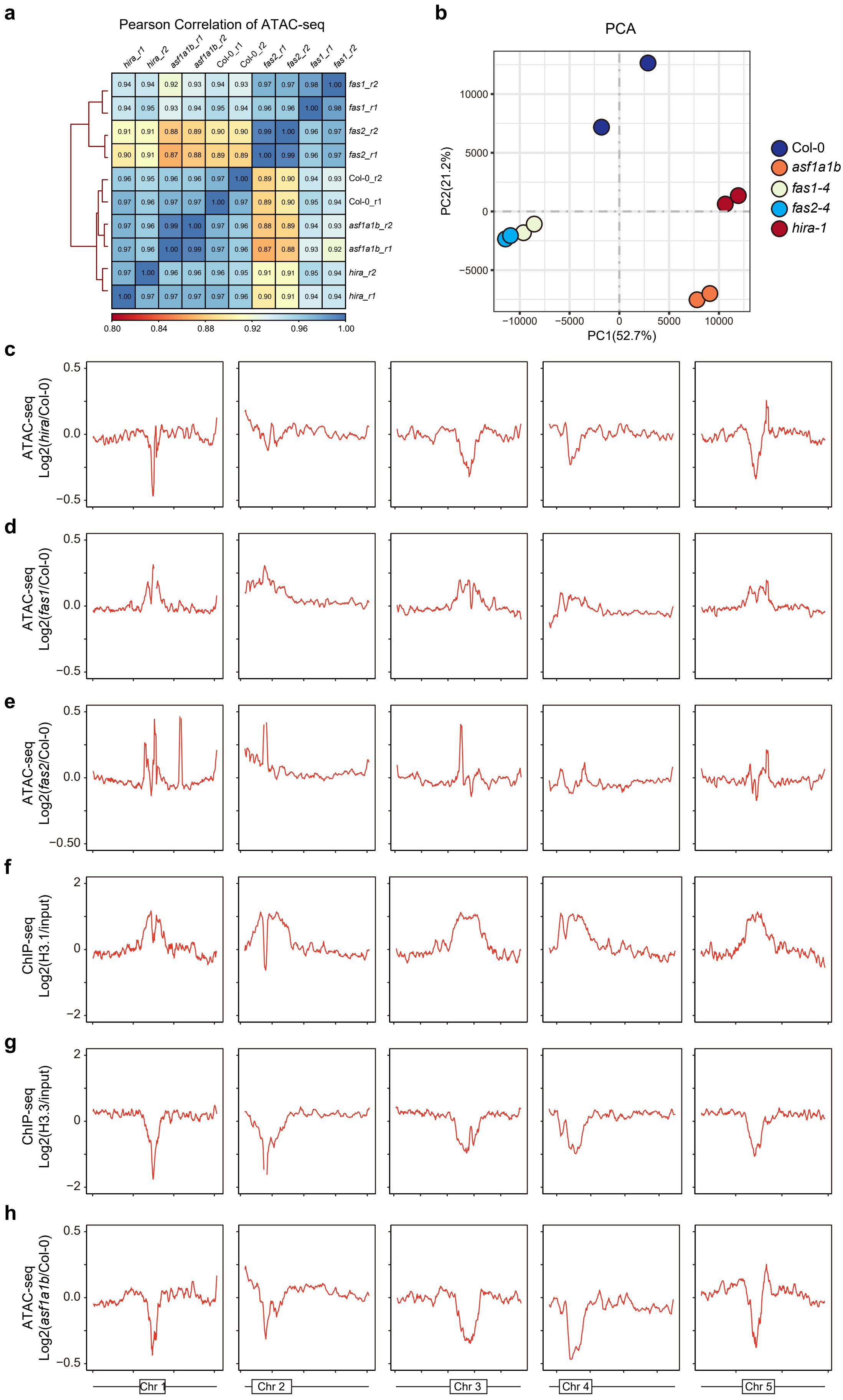
a
Pearson Correlation of ATAC-seq
hira_r1 hira_r2 asf1a1b_r1 asf1a1b_r2 Col-0_r1 Col-0_r2 fas2_r1 fas2_r2 fas1_r1 fas1_r2
0.94 0.94 0.92 0.93 0.94 0.93 0.97 0.97 0.98 1.00 fas1_r2
0.94 0.95 0.93 0.94 0.95 0.94 0.96 0.96 1.00 0.98 fas1_r1
0.91 0.91 0.88 0.89 0.90 0.90 0.99 1.00 0.96 0.97 fas2_r2
0.90 0.91 0.87 0.88 0.89 0.89 1.00 0.99 0.96 0.97 fas2_r1
0.96 0.95 0.96 0.96 0.97 1.00 0.89 0.90 0.94 0.93 Col-0_r2
0.97 0.96 0.97 0.97 1.00 0.97 0.89 0.90 0.95 0.94 Col-0_r1
0.97 0.96 0.99 1.00 0.97 0.96 0.88 0.89 0.94 0.93 asf1a1b_r2
0.97 0.96 1.00 0.99 0.97 0.96 0.87 0.88 0.93 0.92 asf1a1b_r1
0.97 1.00 0.96 0.96 0.96 0.95 0.91 0.91 0.95 0.94 hira_r2
1.00 0.97 0.97 0.97 0.97 0.96 0.90 0.91 0.94 0.94 hira_r1
0.80 0.84 0.88 0.92 0.96 1.00
b
PCA
PC2(21.2%)
PC1(52.7%)
10000
5000
0
−5000
−10000
Col-0
asf1a1b
fas1-4
fas2-4
hira-1
c
ATAC-seq Log2(hira/Col-0)
d
ATAC-seq Log2(fas1/Col-0)
e
ATAC-seq Log2(fas2/Col-0)
−0.50
f
ChIP-seq Log2(H3.1/input)
g
ChIP-seq Log2(H3.3/input)
h
ATAC-seq Log2(asf1a1b/Col-0)
0.5
0.0
−0.5
2
0
−2
Chr 1
Chr 2
Chr 3
Chr 4
Chr 5

**Figure S1. Comparison of ATAC-seq and H3s distribution.**
**(a)** Pearson correlation coefficient between replicates of Col-0, asf1a1b, hira-1, fas1-4, and fas2-4 ATAC-seq data.
**(b)** Principal component analysis (PCA) from replicates of Col-0, asf1a1b, hira-1, fas1-4, and fas2-4 ATAC-seq data.
Differential (mutant vs Col-0) ATAC-seq signals plotted over Arabidopsis chromosomes in hira-1 **(c)**, fas1-4 **(d)**, fas2-4 **(e)** and asf1a1b **(h)**. H3.1 **(f)** and H3.3 **(d)** ChIP signals in Col-0 over Arabidopsis chromosomes. Reads per million mapping reads (RPKM) is calculated in 100-kb bins and normalized with Col-0 or ChIP input. Boxes correspond to the heterochromatic region.

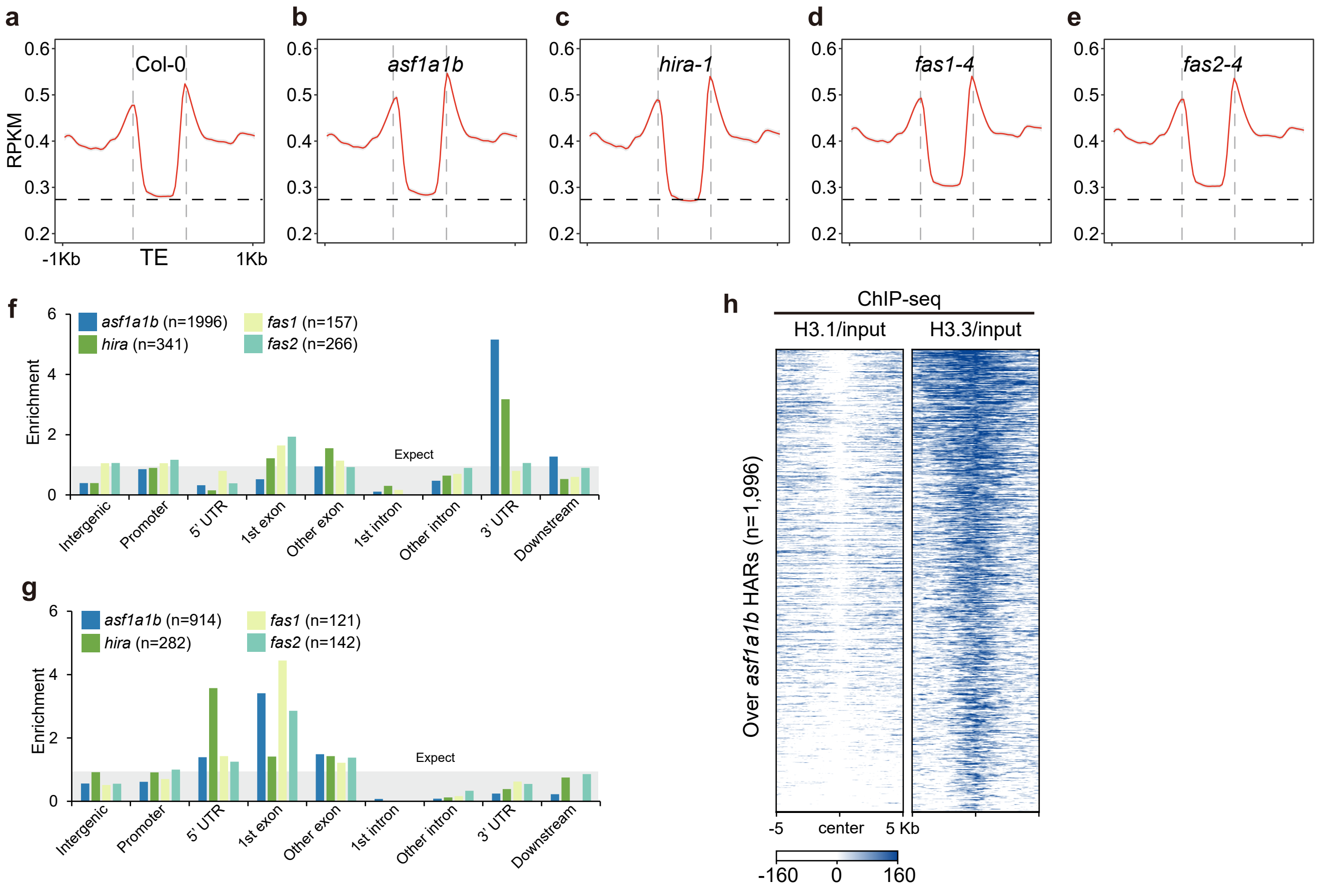


**Figure S2. Distribution of ATAC-seq reads over transposon elements (TE) and genomic distribution of the differentially enriched ATAC-seq peaks.**

Metaplot of Col-0 **(a)**, asf1a1b **(b)**, hira-1 **(c)**, fas1-4 **(d)**, and fas2-4 **(e)** ATAC-seq data over TE (n=32153) including 1 Kb flanking sequence.

**(f)** Genomic distribution of asf1a1b, hira-1, fas1-4, and fas2-4. Highly Accessible Regions (HARs) represent genomic regions where mutants have higher chromatin accessibility than wild-type.

**(g)** Genomic distribution of asf1a1b, hira-1, fas1-4, and fas2-4. Lowly Accessible Regions (LARs) represent genomic regions where mutants have lower chromatin accessibility than wild-type ones. Enrichment = (number of sample peaks)/(number of randomly selected peaks).

**(h)** Heatmap showing ChIP-seq signals of H3.1/input, H3.3/input over asf1a1b HARs (n=1,996).

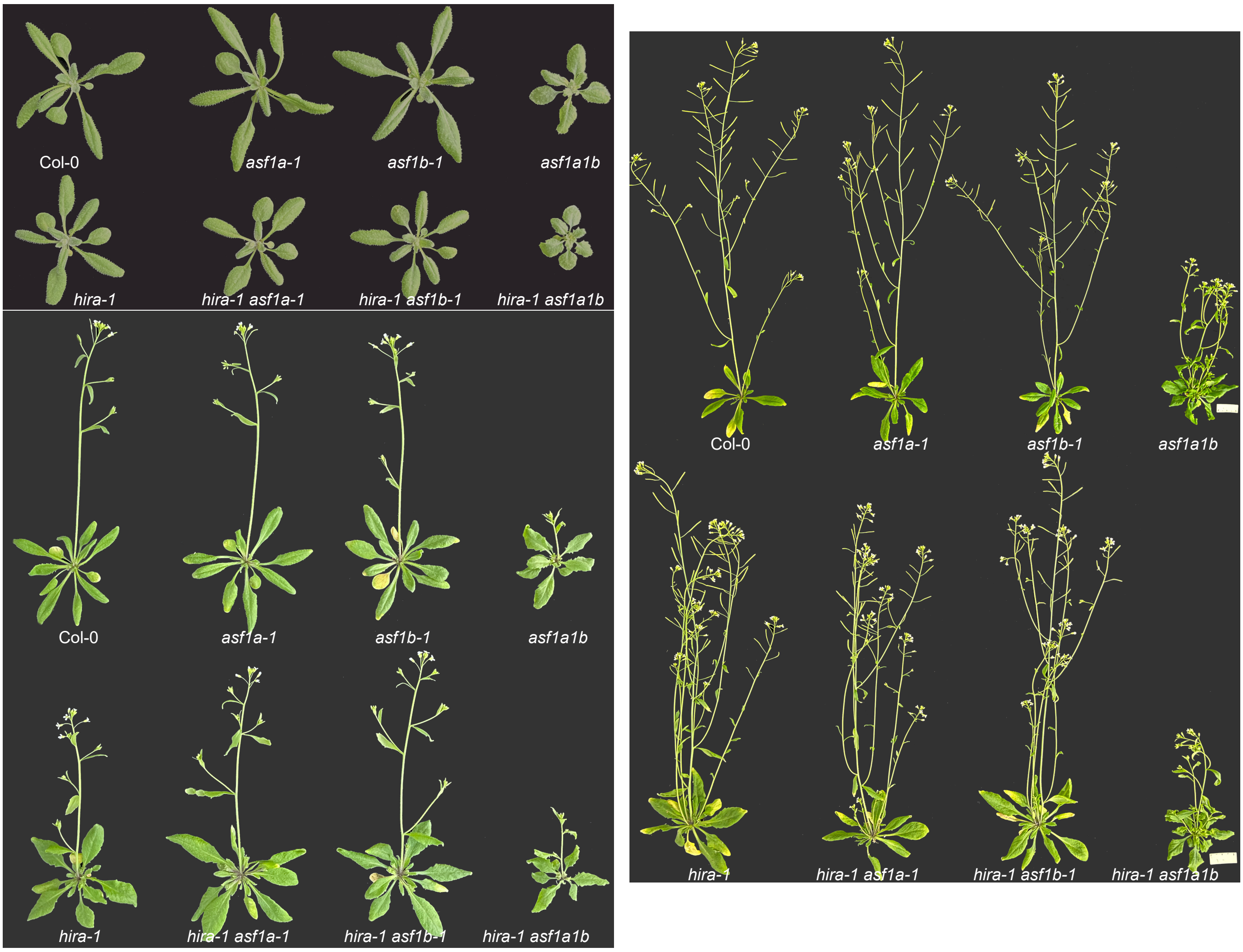


**Figure S3.** Morphology of Col-0, asf1a-1, asf1b-1, hira-1, asf1a1b, hira-1 asf1a-1, hira-1 asf1b-1, and hira-1 asf1a1b.

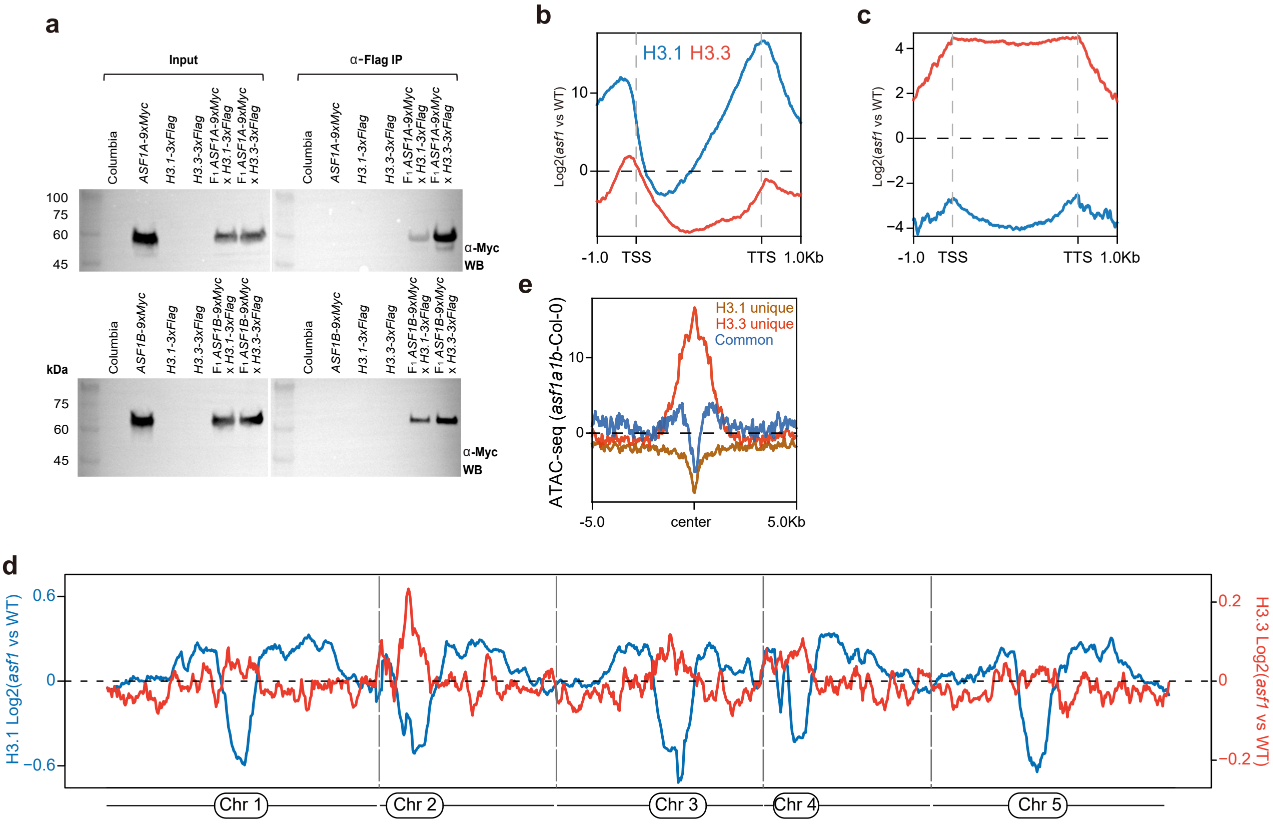


**Figure S4. ASF1 mutation causes a redistribution of H3.3 and H3.1.**
**(a)** Co-immunoprecipitation assay of ASF1A, ASF1B, H3.1, and H3.3. Labels to the right indicate the antibody used for the western blot.
**(b)** Metaplot of H3.1 and H3.3 variation (asf1a1b vs. Col-0) over the gene body and 1 Kb flanking sequence of genes with coding regions lengths over 150bp (n=26816). The Blue and the red line represents H3.1 and H3.3 variation, respectively.
**(c)** Metaplot of H3.1 and H3.3 variation (asf1a1b vs Col-0) over TE (n=32153) including 1 Kb flanking sequence. The Blue and the red line represents H3.1 and H3.3 variation, respectively.
**(d)** Metaplot of H3.1 and H3.3 variation (asf1a1b vs Col-0) over 5 chromosomes splitting into 100 Kb bins. The Blue and the red line represents H3.1 and H3.3 variation, respectively.
**(e)** Variation of ATAC-seq (asf1a1b vs Col-0) over H3.1-unique (n=24841), H3.3-unique (n=28629), and H3.1/H3.3 common (n=15051) peaks.

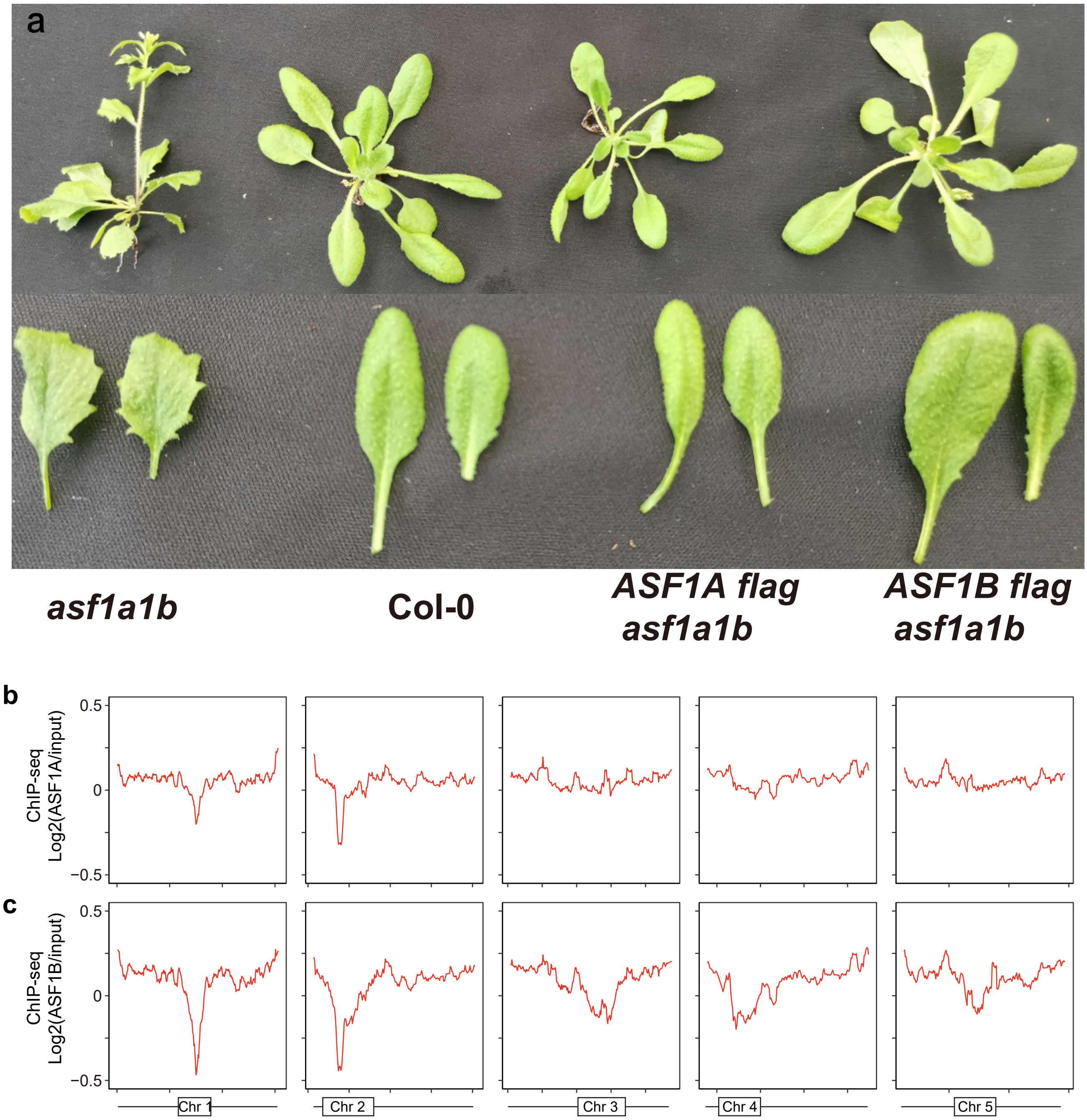


**Figure S5. Genomic distribution of ASF1 proteins.**
**(a)** Morphological phenotype of asf1a1b mutants transformed with ASF1A-Flag and ASF1B-Flag.
**(b-c)** Whole-genome distribution of ASF1A and ASF1B ChIP-seq signal over Arabidopsis chromosomes. Reads per million mapping reads (RPKM) were calculated in 100-Kb bins and normalized with Col-0 or ChIP input. Boxes correspond to the heterochromatic regions.

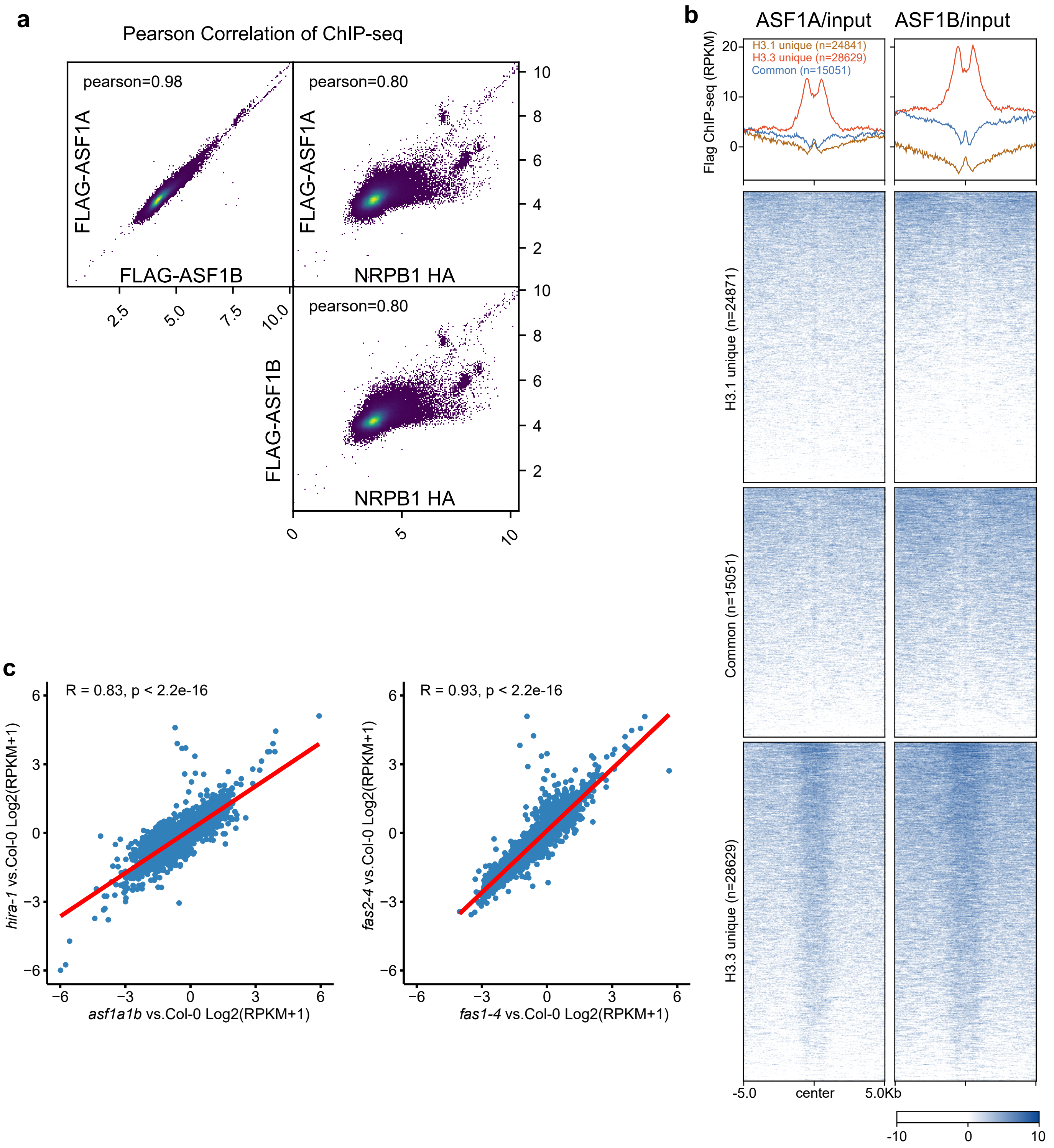


**Figure S6. Correlation between ASF1A, ASF1B, NRPB1 and H3.1/H3.3 ChIP-seqs.**
**(a)** Genome-wide pairwise scatterplots for ChIP-seq at 1 Kb bins.
**(b)** Plotting of ASF1A and B ChIP-seq signal over H3.1 unique (n=24841), H3.3 unique (n=28629), and H3.1/H3.3 common (n=15051) peaks.
**(c)** Correlation of the expression divergence between H3 chaperone mutants. Graph showing correlation of variation in RNA-seq level of genes (n=3412) that are differentially expressed (FC > 2, FDR < 0.05) in at least one mutant in asf1a1b vs hira-1and fas1-4 vs fas2-4.

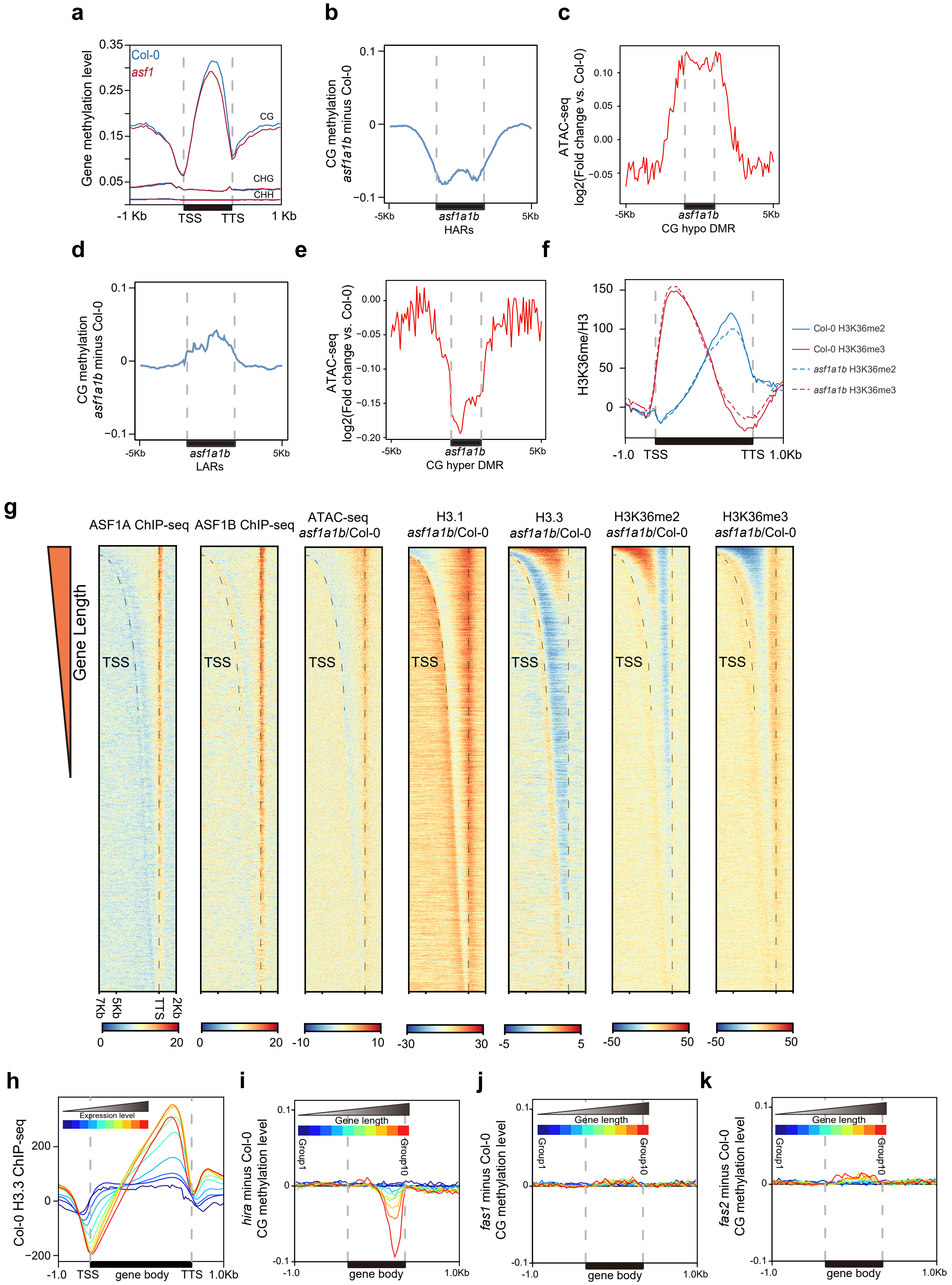
a
0.35
Col-0
asf1
0.25
0.15
0.05
Gene methylation level
CG
CHG
CHH
-1 Kb
TSS
TTS
1 Kb
b
0.1
0
−0.1
CG methylation
asf1a1b minus Col-0
-5Kb
asf1a1b
HARs
5Kb
c
0.10
0.05
0.00
−0.05
ATAC-seq
log2(Fold change vs. Col-0)
-5Kb
asf1a1b
CG hypo DMR
5Kb
d
0.1
0
−0.1
CG methylation
asf1a1b minus Col-0
-5Kb
asf1a1b
LARs
5Kb
e
0.00
−0.05
−0.10
−0.15
−0.20
ATAC-seq
log2(Fold change vs. Col-0)
-5Kb
asf1a1b
CG hyper DMR
5Kb
f
150
100
50
0
H3K36me/H3
-1.0
TSS
TTS
1.0Kb
Col-0 H3K36me2
Col-0 H3K36me3
asf1a1b H3K36me2
asf1a1b H3K36me3
g
Gene Length
ASF1A ChIP-seq
ASF1B ChIP-seq
ATAC-seq
asf1a1b/Col-0
H3.1
asf1a1b/Col-0
H3.3
asf1a1b/Col-0
H3K36me2
asf1a1b/Col-0
H3K36me3
asf1a1b/Col-0
TSS
7Kb
5Kb
TTS
2Kb
0
20
0
20
-10
10
-30
30
-5
5
-50
50
-50
50
h
Expression level
200
0
−200
Col-0 H3.3 ChIP-seq
-1.0
TSS
gene body
TTS
1.0Kb
i
0.1
0
-0.1
Gene length
Group1
Group10
hira minus Col-0
CG methylation level
-1.0
gene body
1.0Kb
j
0.1
0
-0.1
Gene length
Group1
Group10
fas1 minus Col-0
CG methylation level
-1.0
gene body
1.0Kb
k
0.1
0
-0.1
Gene length
Group1
Group10
fas2 minus Col-0
CG methylation level
-1.0
gene body
1.0Kb

**Figure S7. Effects of asf1a1b on the distribution of epigenetic marks.**
**(a)** Metaplot showing DNA methylation level in Col-0 and asf1a1b over the gene body and 1 Kb flanking sequence.
**(b)** Metaplot showing differential CG methylation level (asf1a1b minus Col-0) over asf1a1b highly accessible regions (HAR, n=1,996).
**(c)** Metaplot showing differential of chromatin accessibility (asf1a1b/Col-0) over asf1a1b CG hypomethylated DMRs (n=6197).
**(d)** Metaplot showing differential CG methylation level (asf1a1b minus Col-0) over asf1a1b lowly accessible regions (LAR, n=914).
**(e)** Metaplot showing differential of chromatin accessibility (asf1a1b/Col-0) over asf1a1b CG hypermethylated DMRs (n=3867).
**(f)** Metaplot showing H3K36me2/3 level in Col-0 and asf1a1b over the gene body and 1 Kb flanking sequence.
**(g)** ChIP-seq (ASF1A, ASF1B, H3.1, H3.3, and H3K36me2/3) and ATAC-seq signal over genes ranked by gene length. Top to bottom display genes from long to short.
**(h)** Metaplot showing H3.3 ChIP-seq signal in Col-0 over genes according to expression level (Group 1 to Group 10 represent gene expression from low to high).
**(i-k)** Metaplot of CG methylation difference in hira-1 **(i)**, fas1-4 **(j)**, and fas2-4 **(k)** vs. Col-0 over genes grouped by gene length (Group 1 to Group 10 represent gene length from short to long).

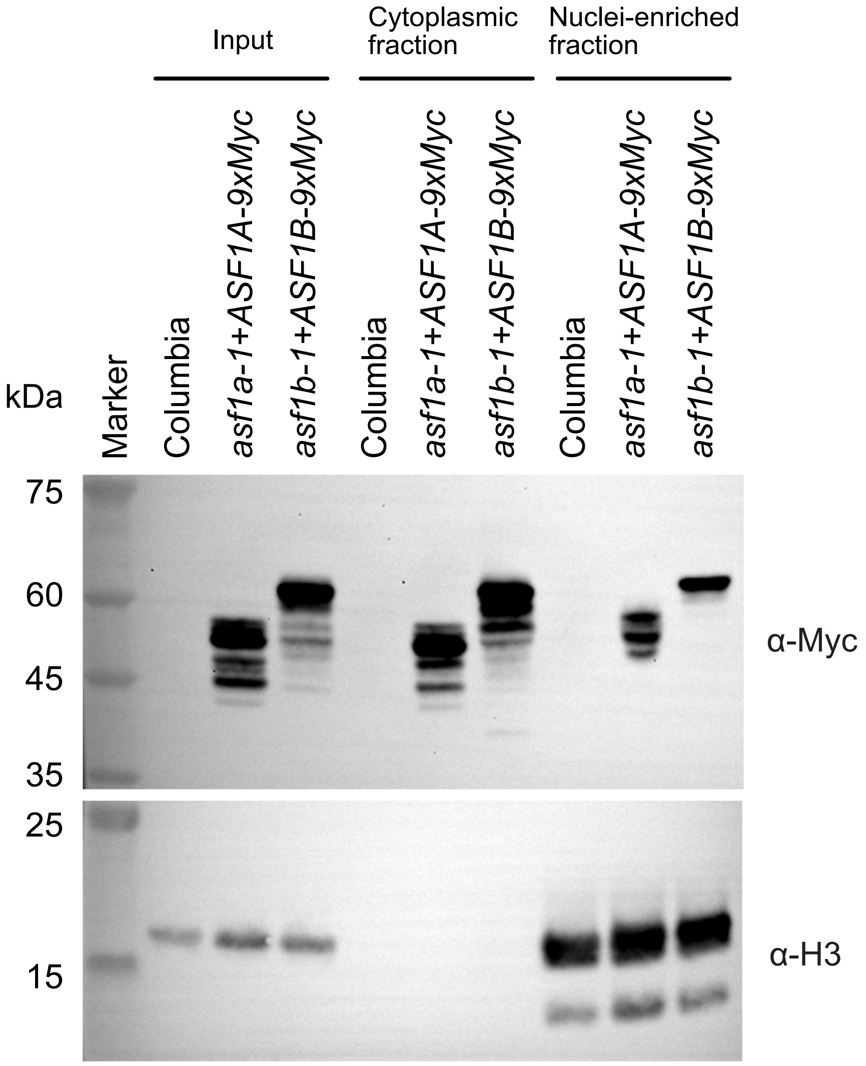


**Figure S8. Subcellular fractionation of ASF1 proteins.**
Western blot analysis of nuclear and cytoplasmic fractions using an anti-Myc antibody for ASF1A and ASF1B proteins. Western blot using anti-H3 antibody is a control for nuclear proteins. Numbers on the left indicate the protein ladders' molecular weights (kDa).

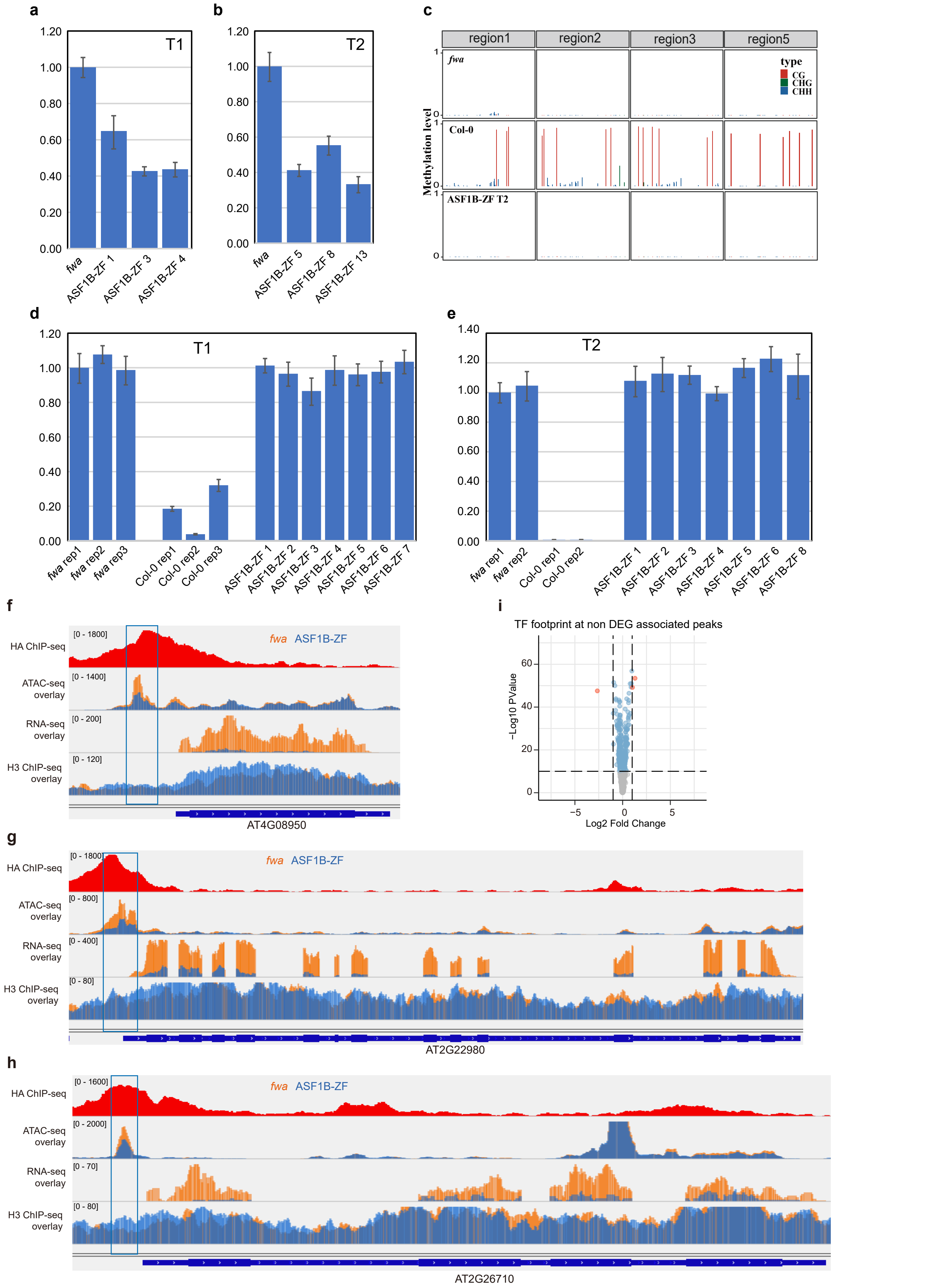


**Figure S9. ASF1B-ZF108 effects on FWA expression and TF footprint.**
**(a-b)** The relative expression level of FWA in fwa and ASF1B-ZF108 T1 lines and T2 lines. **(c)** DNA methylation (CG, CHG, and CHH) level over the FWA promoter using bisulfite (BS)-PCR-seq in Col-0 and fwa-4 controls and the ASF1B-ZF108 T2 line. Light red boxes indicate the ASF1B-ZF108 binding sites.
**(d-e)** McrBC assay of Col-0 and fwa-4 controls and the ASF1B-ZF108 T1 and T2 lines at the FWA tandem repeat.
**(f-h)** Screenshot of example genes showing ZF binding assayed by ChIP-seq, chromatin accessibility assayed by ATAC-Seq, and expression level at AT4G08950, AT2G22980, and AT2G26710. Tracks of fwa-4 (orange) and ASF1B-ZF108-3xFlag (blue) are shown overlapped.
**(i)** Volcano plot showing differential footprint of 572 plant TF motifs distributed at non-DEG associated, down-regulated off-target sites (n=572).
